# COVID-19 infection enhances susceptibility to oxidative-stress induced parkinsonism

**DOI:** 10.1101/2022.02.02.478719

**Authors:** Richard J Smeyne, Jeffrey Eells, Debotri Chatterjee, Matthew Byrne, Shaw M. Akula, Srinivas Sriramula, Dorcas P. O’Rourke, Peter Schmidt

**Author notes:** **<u>corresponding author</u>** Richard Smeyne, Ph.D., Department of Neurosciences, Thomas Jefferson University, Vickie and Jack Farber Institute for Neuroscience, JHN 451, 900 Walnut Street, Philadelphia, PA, 19027. The first three authors contributed equally to this paper. **<u>Complete Author contact information</u>** Richard Smeyne, Ph.D., Jeffery Eels, Ph.D., Debotri Chatterjee, B.A., Matthew Byrne, B.S, Shaw M. Akula, Ph.D, Srinivas Sriramula, Ph.D, Dorcas P. O’Rourke, D.V.M, Peter Schmidt, Ph.D.

## Abstract

**Background:** Viral induction of neurological syndromes has been a concern since parkinsonian-like features were observed in patients diagnosed with encephalitis lethargica subsequent to the 1918 influenza pandemic. Given the similarities in the systemic responses following SARS-CoV-2 infection with those observed after pandemic influenza, there is a question if a similar syndrome of post-encephalic parkinsonism could follow COVID-19 infection.

**Objectives:** To determine if prior infection with SARS-CoV-2 increased sensitivity to a mitochondrial toxin known to induce parkinsonism.

**Methods:** hACE2 mice were infected with SARS-CoV-2 to induce mild to moderate disease. After 31 days recovery, mice were administered a non-lesion inducing dose of the parkinsonian toxin MPTP. Subsequent neuroinflammation and SNpc dopaminergic neuron loss was determined and compared to SARS-CoV-2 or MPTP alone.

**Results:** hACE2 mice infected with SARS-CoV-2 or MPTP showed no SNpc DA neuron loss following MPTP. In mice infected and recovered from SARS-CoV-2 infection, MPTP induced a 23% or 19% greater loss of SNpc dopaminergic neurons than SARS-CoV-2 or MPTP, respectively (*p*□<□0.05).

Examination of microglial activation showed a significant increase in the number of activated microglia in the SARS-CoV-2 + MPTP group compared to SARS-CoV-2 or MPTP alone.

**Conclusions:** Our observations have important implications for long-term public health, given the number of people that have survived SARS-CoV-2 infection as well as for future public policy regarding infection mitigation. However, it will be critical to determine if other agents known to increase risk of PD also have synergistic effects with SARS-CoV-2 and if are abrogated by vaccination.

**Funding:** This work was supported by grant from the State of North Carolina (PS, JE, DOR, RJS) and R21 NS122280 (RJS).

## Introduction

Neurological sequalae subsequent to viral infection have been reported since von Economo’s description of a post-influenza encephalopathy, including aspects of parkinsonism that were reported to the Vienna neurological society in 1917 ^1^. These observations were consistent with the post-encephalic symptoms that followed the 1918 H1N1 (Spanish) influenza pandemic and persisted for about 10 years subsequent to its zenith ^2, 3^. While the 1918 H1N1 influenza was thought to be particularly virulent and its sequalae particularly devastating, it has not been the only viral outbreak linked to post-encephalic symptoms ^4^. One of the more common symptoms of these viral encephalopathies is the development of parkinsonism ^4^. The mechanism for this is reportedly linked to viral affinity for the highly vascularized midbrain catecholaminergic neurons in the substantia nigra and locus coeruleus ^5^ that are lost in Parkinson’s disease. Mechanisms of indirect action after viral infection, such as effects of inflammatory cytokines or glia activation have been demonstrated previously ^6, 7^.

In 2019, a novel coronavirus outbreak was reported in China and in the ensuing pandemic, nearly 300 million cases have been reported worldwide ^8^. The virus, SARS-CoV-2, is a large enveloped non segmented positive sense RNA virus ^9^. Like its related family members SARS-CoV and MERS, it originally presents as predominantly a respiratory illness ^10^; however, a number of other organ systems ^11^, including the nervous system, are also severely affected ^12^. Relating specifically to SARS-CoV-2, it is unclear if these effects are direct, based on the virus’ ability to enter the brain ^13^, or if they arise via a peripheral mechanism such as an induction of a cytokine storm that induces neurological changes in the peripheral immune system which then transmits its signals to the brain ^14^.

Given the long history of viral infections inducing basal ganglia disease ^4^ combined with the scale and scope of the COVID-19 pandemic, it is incumbent to explore if there might be an increased risk of similar neurological sequelae among individuals recovered from mild to moderate or severe COVID-19. Recent reports have empirically associated SARS-CoV-2 infection with development of clinical parkinsonism ^15-17^. In order to experimentally test the hypothesis that prior infection with SARS-CoV-2 could directly increase the risk of parkinsonian pathology through viral mechanisms identified previously, we evaluated nigrostriatal degeneration in a preclinical mouse model where genetically-tailored susceptible animals, engineered to express the human ACE2 receptor ^18^, were infected with SARS-CoV-2 (strain USA-1) and allowed to recover. Thirty-one days after the animals recovered from the viral infection, we challenged them with subtoxic levels of a mitochondrial toxin, 1-methyl-4-phenyl-1,2,3,6-tetrahydropyridine (MPTP); a chemical that is known to block complex I and IV of the electron transport chain and induce some of the characteristic pathologies seen in Parkinson’s disease^19^ We found that animals that recovered from SARS-CoV-2 infection were more susceptible to the parkinsonian effects of non-lethal levels of MPTP than mice infected with SARS-CoV-2 or administered MPTP alone. This preclinical study suggests that infection with SARS-CoV-2 is a likely predisposing risk factor for later development of Parkinson’s disease.

## Methods

All the work pertaining to the use of SARS-CoV-2 were performed in BSL-3 laboratory. SARS-CoV-2 (isolate USA-WA1/2020), obtained from bei RESOURCES (Manassas, VA), was propagated in Vero cells using a 10-30% sucrose gradient in an ultracentrifuge (Beckman L8-55), and the yield titrated using the classic Reed and Muench formula ^20^. Viral studies were approved by the Office of Prospective Health/Biological Safety for the use of biohazardous agent (SARS-CoV-2) and the registration number is 20-01 (title: Host response to COVID-19 infection in Eastern North Carolina).

To evaluate the risk neurological sequelae among individuals recovered from moderate or severe COVID-19, we designed a study to analyze established mechanistic pathways for viral induction of basal ganglia disease, not previously tested with SARS-CoV2 infection. Working under an IACUC-approved protocol (AUP#A209) in a fully AAALAC-accredited facility, SARS-CoV2 susceptible mice (B6.Cg-Tg(K18-ACE2)2Prlmn/J, Jackson Labs, Strain #034860, hACE2 mice) were randomly assigned to a dose-finding study and then four study arms. First, in a dose-finding study, mice were tested at three doses of virus (three each at 10^3^, 4×10^3^, and 10^4^ TCID_50_) and the optimal dose was selected. Eighteen mice in total were infected intranasally with SARS-CoV-2 at the optimal study dose (strain USA-1 obtained from BEI Resources)), dose 4×10^3^ TCID_50_ in 25 ml saline divided equally into each naris) and eleven were subjected to a sham procedure (25μl saline total, 12.5μl into each naris). The singly-housed mice were managed through their infection and recovery in an ABSL3 facility (East Carolina University, Greenville, NC). Thirty-one days after the animals recovered from the viral infection, six recovered SARS-CoV2 and five sham-treated mice were challenged with MPTP, a mitochondrial stressor known to induce some of the characteristic pathologies seen in Parkinson’s disease; and has been previously used in similar studies ^21^. We used a subtoxic dosage of MPTP (10mg/kg x 4 at 2 hour intervals, injected ip) that has been shown to induce a small inflammatory response, but no dopamine neuron death^21^. The remaining animals were injected ip with equal volumes of saline using the same procedure. For safety reasons, mice were anesthetized with 3% isoflurane prior to each injection. Animals were observed for signs of infection, decreases in body weight, alterations in body temperature, lack of grooming, hunched posture, rapid respiration, lethargy, and mortality and were managed under the supervision of an expert research veterinarian (author O’Rourke). Continuous monitoring of animal temperature and activity was provided by an RFID microchip and sensors using the UID Mouse Matrix and UID Temperature Microchips from uidevices (www.uidevices.com). To confirm infection in mice, blood was collected via cardiac puncture prior to perfusion and placed in KE EDTA tubes and plasma was isolated. Antibody titers were measured in plasma using the Mouse Anti-SARS-CoV-2 IgG Antibody ELISA Kit (DEIASL240) from CD Creative Diagnostics (www.creative-diagnostics.com) according to manufacture instructions.

Forty-five days after active or sham infection and 14 days after MPTP or saline administration, the animals were anesthetized and euthanized by transcardial perfusion with 3% paraformaldehyde. Brains were removed from the calvaria and postfixed in fresh fixative. Brain were processed, embedded and sectioned for histopathological evaluation. SNpc dopamine neuron number, using immunohistochemical detection of tyrosine hydroxylase and counterstaining with a Nissl stain, and analysis of total, resting and activated microglia was performed using stereological methods as described ^22^.

## Results

### Effects of SARS-CoV-2 infection on animal morbidity and mortality

B6.Cg-Tg(K18-ACE2)2Prlmn/J mice ((hACE2 mice) were infected intranasally with one of 3 different titers (1 × 10^3^ TCID_50_, 4 × 10^3^ TCID_50_ and 1 × 10^4^ TCID_50_) of SARS-CoV-2. After exposure, the singly-housed mice were examined for signs of infection including a decrease in body weight, alterations in body temperature, lack of grooming, hunched posture, rapid respiration and lethargy. We observed no animal morbidity or mortality in mice infected with 1 × 10^3^ TCID_50_, about 28% in mice infected with 4 × 10^3^ TCID_50_, and 70% in animals infected with 1 × 10^4^ TCID_50_ (**Fig 1A**). The animals that died were found deceased in their cage or were euthanized at a time when they had experienced >20% weight loss and were moribund; a condition usually detected within 24 hours of infection), were hypothermic and appeared to have milky white discharge from their eyes. Animals that survived showed no apparent temperature alterations nor did they show any loss of blood oxygenation. To ensure that surviving mice in this group were infected with the virus, we performed antibody titers 45 days after infection. As seen in **Fig 1B**, there was a significant antibody response in both groups (naïve and MPTP-treated) infected with SARS-CoV-2. No detectable viral titer was measured in animals intranasally administered the saline alone. Based on this empirical data, we examined the effects of subtoxic levels of MPTP on inflammation and dopaminergic neuron death in the animals infected with the moderate dose.

**Figure 1.**
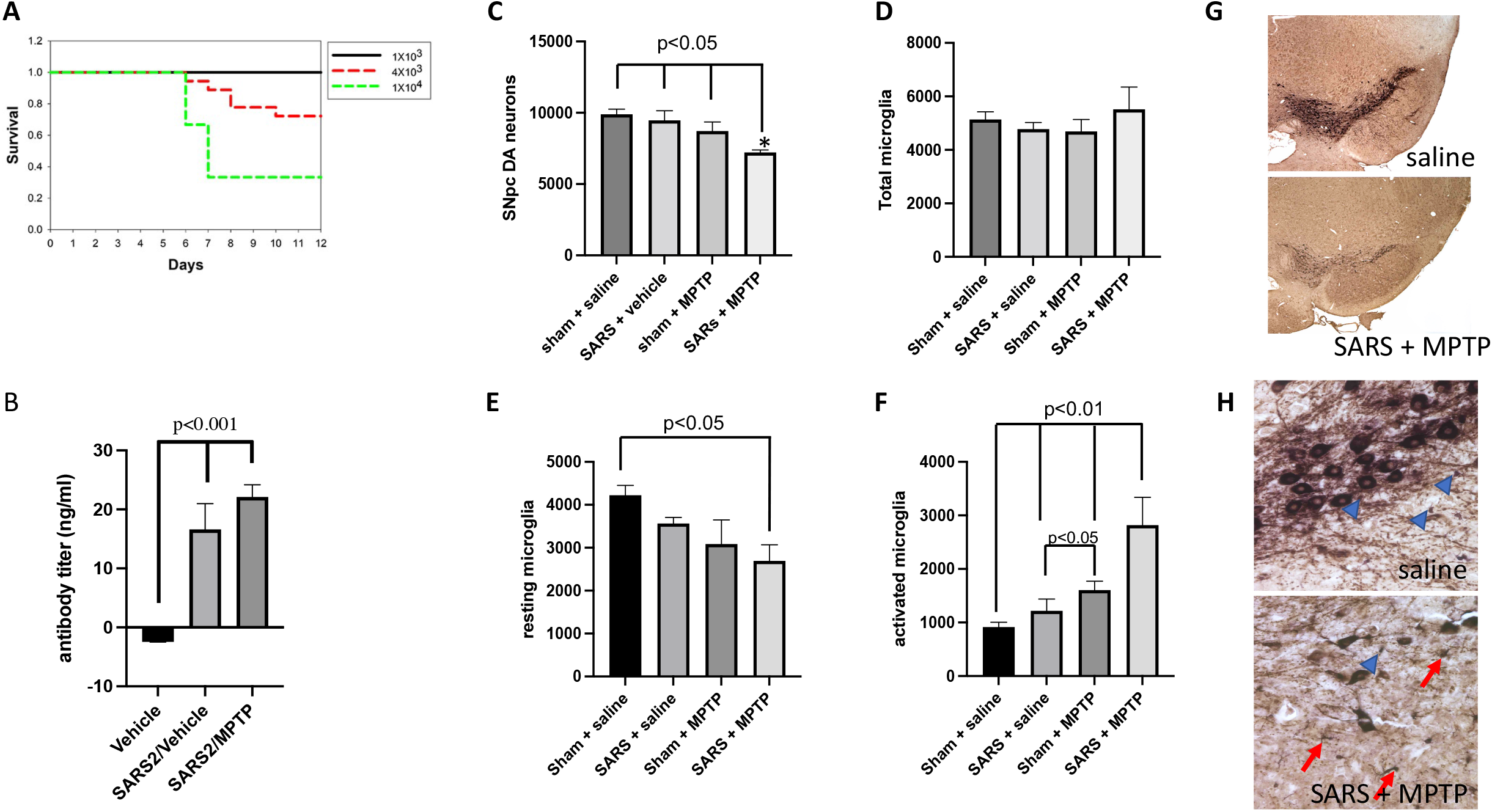
Effect of SARS-CoV-2 infection on sensitivity to subacute MPTP infection. A. Kaplan-Meier Survival curve of mice infected with 3 different titers of SARS-CoV-2 (USA-1). B. Antibody titers in mice infected 45 days prior to testing. C. Number of SNpc dopaminergic neurons in the SNpc. D. Total number of microglia in the SNpc. E. Total number of resting microglia in the SNpc. F. Total number of activated microglia in the SNpc. G. Low power (4x) micrograph showing appearance of the dopaminergic neurons (TH+ neurons) in the rostral tier of the substantia nigra pars compacta. The SARS + MPTP group has a significant loss of these neurons as well as its fibers as noted by the decreased TH-immunoreactivity. H. Higher power photomicrograph (40x) of the same sections shown in G. Resting microglia are small and have thin processes and are labeled with blue arrowheads. Activated microglia have a larger cell body, and thickened processes; labeled by red arrows. Statistical analysis was performed using ANOVA with post-hoc tests (Tukey) (Prism 9.0, GraphPad Software) if overall significance was achieved.

### Prior infection with SARS-CoV-2 increases susceptibility to the dopaminergic toxin, MPTP

In this study, we administered a subtoxic dosage of MPTP (10mg/kg x 4, ip injections every 2 hours) that we have previously and empirically shown to induce a small inflammatory response but no dopamine neuron death ^21^. We examined the effects of these low levels of MPTP on 4 groups of mice: 1) hACE2 mice intranasally administered saline, 2) hACE2 mice intranasally administered SARS-CoV-2, 3) hACE2 mice intranasally administered saline and 31 days later given MPTP and 4) hACE2 mice intranasally administered SARS-CoV-2 and after 31 days given MPTP. We then waited 14 days after MPTP and killed animals to stereologically assess both SNpc DA neuron and microglia numbers. As seen in **Fig 1C**, no SNpc DA neuron loss was observed after saline, SARS-CoV-2 alone or MPTP alone. However, in mice infected with 4 × 10^3^ TCID_50_ SARS-CoV-2 and allowed to recover for 30 days, this concentration (10mg/kg MPTP x 4) induced a significant 23% or 19% greater loss of substantia nigra pars compacta dopaminergic neurons than SARS-CoV-2 or MPTP, respectively (**Fig 1C, G**, *p*□<□0.05).

### Prior infection with SARS-CoV-2 increases neuroinflammation

To examine if SARS-CoV-2 infection induced a neuroinflammatory response, we performed a quantitative stereological analysis of total, resting and activated microglia in the SNpc 45 days after SARS-CoV-2 infection; with or without 10mg/kg x 4 MPTP. Infection with SARS-CoV-2 resulted in no change in the total number of microglia in any of the 4 experimental groups (**Fig 1D**). However, when we quantified resting vs activated microglia individually, we observed significant differences between groups. Resting microglia numbers trended lower 14 days following SARS-CoV-2 infection or MPTP alone. In the SARS-CoV-2 + MPTP group, we observed a significant 36% reduction in resting microglia number compared to the saline group (p<0.05). Quantitation of active microglia showed that SARS-CoV-2 infection alone, compared to saline treatment, did not induce an increase in their number. Although the sublethal dose of MPTP did not induce SNpc DA neuron loss (**Fig 1C**), it did induce a 76% (p<0.01) or 43% (p<0.05) increase in the number of activated microglia (**Fig 1D**, p<0.05) compared to saline or SARS-CoV-2 respectively. Mice treated with MPTP after recovering from SARS-CoV-2 infection had a significantly larger 3-fold increase in activated microglia (p<0.01) compared to MPTP alone **(Fig 1F,H)**.

## Discussion

Through 2021, approximately 300 million people worldwide have been infected with SARS-CoV-2; of which fewer than 2% have been reported to have died from the disease ^8^. The majority of these cases involve the original strain isolated in 2020 (USA-1, aka alpha; strain B.1.1.7), although the recent variants delta (strain B.1.617.2) and omicron (strain B.1.1.529), may soon become the dominant infective strain(s) due to their increased transmissibility ^23^. While the predominant symptom that is associated with this virus is respiratory, it has also been shown that a significant proportion, estimated at 35-40% also demonstrate neurological sequalae ^24^ including anosmia, headache, seizures, stroke (hemorrhagic and thrombosis), meningitis and Acute Disseminated Encephalomyelitis (ADAM). Additionally, a subset of these infected individuals manifest neurological symptoms that appear to have a protracted course, i.e “long-haulers”. These individuals describe issues related to confusion or “brain fog”, persistent headache, numbness/tingling, loss of sense of small/taste, dizziness, pain, and blurred vision ^25^. Related to longer-term issues in the post-COVID-19 infection period, one also needs to remain cognizant of the possibility or the development of post-encephalic syndromes that are known to occur following pandemic viral outbreaks ^26^. Perhaps the most famous of these is the development of a both an immediate as well as post-encephalic parkinsonism that occurred subsequent to the 1918 influenza ^3, 27^.

Given recent reports of COVID-19-induced parkinsonism ^15-17^, where the parkinsonian symptoms appear similar to those described after the 1918 H1N1 outbreak, it would be derelict to not consider that these two vastly different pandemic viruses may share a common mechanism that leads to these neurological sequalae. Both influenza and SARS-CoV-2 are respiratory viruses, both have been shown to infect epithelial cells in the lung and gut ^28^ and both the 1918 H1N1 influenza virus ^29^ and SARS-CoV-2 ^30^ do not appear to have significant neurotropic potential (except in cases where there is a breach in the blood-brain barrier). Related to a possible mechanism, both the 1918 influenza and SARS-CoV-2 appear to induce an enhanced program of induction of pro-inflammatory cytokines and chemokines, known as a cytokine storm^31^. These circulating peripheral cytokines and chemokines can easily penetrate the blood-brain barrier through capillary beds as well as communicate with brain parenchyma through brain glymphatics^32^. Once these inflammatory proteins are in the brain, they have been shown to activate the innate immune system of the brain, (microglia and astrocytes) which also begin to secrete inflammatory proteins that have been shown to sensitize neurons to later insults ^33^.

The cellular composition of the substantia nigra pars compacta (SNpc), which is the main CNS region that degenerates in Parkinson’s disease, is unique in that it contains the highest ratio of microglia:neurons within the CNS. This skewed microglia:neuron ratio places the SNpc at a higher risk for reactive oxygen induced damage^34^ and disruption of mitochondrial function ^35^. This cellular damage does not necessarily have to lead to an immediate effect, but can lead to the neurons having a long-term diminished capacity to handle insults. The aforementioned “hit and run” effect would then lead cells to have a lower threshold for survival following future insults that could include any other agent/environmental ^36^ exposure or even genetic sensitivity ^37^ that has been shown to be associated with Parkinson’s disease.

In this study, we modeled in mice genetically modified to express the ACE2 receptor necessary for infection with SARS-CoV-2, the combination of recovery from a moderate COVID-19 infection with later subthreshold mitochondrial inhibition. Our findings that SARS-CoV-2 infection, alone, did not induce CNS inflammation nor SNpc neuron death suggests that this virus, without any other pathology that reduces blood brain barrier break, is not a direct parkinsonian agent. However, we do find that systemic infection appears to sensitize the SNpc dopaminergic neurons to mitochondrial stress that, in and of itself, does not induce neuron loss. This sensitization appears to remain for a period of time after resolution of the viral infection and without any apparent physical manifestation of a direct viral inflammatory effect in the SNpc. This post-infection sensitization of the SNpc dopaminergic neurons is similar to previous studies that have looked at other viruses associated with post-encephalic parkinsonism including the H1N1 and H5N1 influenza viruses.

While these preclinical models of infection leading to later parkinsonism are critical to our understanding of potential downstream outcomes of pandemics, one must also validate the use of our preclinical model. Studies that have examined the cytokine responses in hACE2 mice and humans following infection with SARS-CoV-2 shows a similar cell-type infectivity as well as a similar induction of cytokine/chemokines ^38^. In both mice and humans, the period of infectivity also appears similar. Additionally, using a similar approach to this study, we previously showed that mice that recovered from an influenza (strain ca/09 H1N1) infection developed enhanced susceptibility to the parkinsonian toxin MPTP ^39^. The role of H1N1 as a susceptibility agent has been recently validated in a retrospective study examining risk for developing Parkinson’s disease in humans following influenza^40^. This study showed that previous influenza infection resulted in the risk for developing Parkinson’s disease increased by 173% compared to individuals not infected ^40^. This increase susceptibility was within the confidence interval determined epidemiologically for people born during the time of the 1918 H1N1 pandemic ^41^. Our preclinical studies examining SARS-CoV-2 infection suggest the possibility of a similar transient increase in parkinsonian incidence. Should this risk manifest, the diverse consequences would represent a great burden on patients, families, and society.

Understanding this risk should be a priority. Evaluation of the sensitivity of this mechanism to viral load, and heterogeneity across variants are both important. While diverse environmental agents have been associated with Parkinson’s risk, characterizing the effects of the “second hit” across the range of environmental agents ^42, 43^ beyond the mitochondrial Complex I and IV inhibitor used here will be necessary. We also need to see if any of the current treatments for COVID-19 infection can moderate this viral sensitization. It has been shown that prior vaccination against H1N1 or immediate treatment with Oseltamivir phosphate (Tamiflu™) can eliminate the H1N1 and MPTP synergy ^39^. Thus, vaccination and antiviral therapies have the potential to modify this mechanism. However, for the more than 100 million people worldwide who survived COVID-19 without the benefit of access to vaccinations, the long-term consequences of infection, including increasing the risk for developing Parkinson’s disease, need to be understood. It is also critical for our healthcare providers and governmental agencies to prepare for this potential.

## Acknowledgements

The authors thank Morgan Alston (TJU) for assistance with histology and John F. Williams, Jason St. Antoine and Matthew Verzwyvelt (ECU) for laboratory assistance in the BSL3 facility. We also thank Michelle Smeyne, Elena Kozina and Aliya Yu for critical reading of the manuscript.

## Author’s Roles

RJS: designed study, performed experiments, analyzed data, wrote paper

JE: designed experiments, performed experiments, analyzed data

DC: performed experiments, analyzed data

DPO: performed experiments, analyzed data

MB, performed experiments

SMA: performed experiments

SS: performed experiments

PS: designed study, analyzed data, wrote paper, arranged funding for study.

## Financial Disclosures of all authors

***Stock Ownership in medically-related fields***: None

***Intellectual Property Rights***: None

***Consultancies***: DOR: Council member emeritus for AAALAC International.

***Expert Testimony***: None

***Advisory Boards***: RJS: Chair of the Scientific Advisory Board of the Parkinson’s Foundation; DOR: North Carolina Association for Biomedical Research (BOD), Sylvan Heights Waterfowl Park (BOD); SA: Scientific Reports;

***Employment***: RJS, DC, MD: Thomas Jefferson University, JE, SA, SS, DOR, PS: East Carolina University.

***Partnerships***: None ***Inventions***: None ***Contracts***: None

***Honoraria***: DOR: AAALAC

***Royalties***: None

***Patents***: Cells that lack p19INK4D and p27KIP1 activity and methods of use thereof. United States Patent 6589505 July 8, 2003; Method for Detection of Adenosine and Metabolite Thereof, US Patent 9528976 12/27/2016 (RJS)

***Grants*** NIH: R01NS110084, R21NS122280, NIEHS:R01ES030937 (RJS), R01HL153115 (SS), State of North Carolina (RJS, JE, SA,SS,DOR PS)

***Other:*** *None*

